# Sex differences in the reward value of familiar mates in prairie voles

**DOI:** 10.1101/2021.09.01.458610

**Authors:** Daniel M. Vahaba, Emily R. Halstead, Zoe R. Donaldson, Todd H. Ahern, Annaliese K. Beery

## Abstract

The rewarding properties of social interactions facilitate relationship formation and maintenance. Prairie voles are one of the few laboratory species that form selective relationships, manifested as “partner preferences” for familiar partners versus strangers. While both sexes exhibit strong partner preferences, this similarity in outward behavior likely results from sex-specific neurobiological mechanisms. We recently used operant conditioning to demonstrate that females work harder for access to a familiar versus unfamiliar conspecific of either sex, while males worked equally hard for access to any female, indicating a key sex difference in social motivation. As tests were performed with one social target at a time, males might have experienced a ceiling effect, and familiar females might be more relatively rewarding in a choice scenario. Here we performed a social choice operant task in which voles could repeatedly lever-press to gain temporary access to either the chamber containing their mate or one containing a novel opposite-sex vole. Females worked hardest to access their mate, while males pressed at similar rates for either female. Individual male behavior was heterogeneous, congruent with multiple mating strategies in the wild. Voles exhibited preferences for favorable over unfavorable environments in a non-social operant task, indicating that lack of social preference does not reflect lack of discrimination between chambers. Oxytocin receptor genotype at the intronic SNP NT213739 replicated a prior association with stranger-directed aggression within the test. These findings suggest that convergent preference behavior in male and female voles results from sex-divergent pathways, particularly in the realm of social motivation.

## Introduction

Prairie voles are socially monogamous rodents that often form lifelong opposite-sex pair bonds in natural environments, and both males and females exhibit selective preferences for familiar mates in the laboratory (Getz *et al*. 1981; Williams *et al*. 1992; Scribner *et al*. 2020). Despite similar partner preference behavior, there are several indications that mechanisms underlying bond formation in males and females may differ—including in the timing of bond formation, reliance on specific neuropeptide signaling pathways, and the role of behavioral reward (De Vries 2004; DeVries & Carter 2011; Beery *et al*. 2021; Brusman *et al*. submitted to this issue). Our lab recently demonstrated that female prairie voles work harder to gain access to mates versus stranger males, whereas males work equally hard to access mates and unfamiliar females (Beery *et al*. 2021), indicating high salience of any female. However, such testing was conducted with only one available social stimulus at a time, leaving open the possibility that males would still work harder for a known mate over an unfamiliar stranger if given an opportunity to do so. In the present study, we developed an operant choice setup to determine whether prairie voles would consistently learn to press for one stimulus vs. another (non-social choice testing), then asked whether males and females would both exhibit preferences for familiar mates versus unfamiliar opposite-sex strangers in operant social choice tests. We then related oxytocin receptor genotype data to affiliative and aggressive behaviors recorded during these tests.

The partner preference test (PPT), developed in the laboratory of Sue Carter, has become the standard method to assess preferences for familiar mates in voles (Williams *et al*. 1992). During the PPT, the focal vole is placed in the middle of a three-chamber apparatus where they can freely explore and spend time near either a familiar vole (e.g. partner) or an unfamiliar vole (e.g. novel, opposite-sex subject/stranger), both of whom are tethered at opposite ends (Williams *et al*. 1992; Beery 2021). Partner preference is assessed by measuring how much time a focal vole spends huddling with each of the stimulus voles. While this method can indicate selective preference for familiar mates, such preferences could emerge for a variety of reasons, such as enhanced motivation to interact with a mate, increased tolerance of a mate, and/or aversion of unfamiliar “stranger” voles. Because subjects can freely interact with stimulus voles, data from a PPT cannot distinguish between these hypotheses.

Social motivation and reward signaling are important components of prairie vole mate relationships. Dopamine signaling is necessary for social bond formation and maintenance between mates (Aragona *et al*. 2003, 2006), although this does not appear to be the case for same-sex peer relationships (Lee & Beery 2021). Opioid signaling is also necessary for partnership maintenance, and interacts with dopamine signaling within the nucleus accumbens (Resendez *et al*. 2012, 2016). Behaviorally, prairie voles show a conditioned preference for cues associated with mates in socially conditioned place preference tests (Ulloa *et al*. 2018; Goodwin *et al*. 2019), although unfamiliar opposite-sex conditioning has not been tested. These studies strongly suggest that social motivation – reflecting a drive to seek a socially rewarding partner – is involved in prairie vole mate relationships.

Previous operant paradigms assessing social motivation in rodents have also relied on focal subjects having only one social option at a time (reviewed in Trezza *et al*. 2011). For example, rates of pressing for pup delivery differ in dams with different lesions (Lee *et al*. 1999), and rates of nose-pokes for access to aggress upon a subordinate mouse differ by drug treatment (Fish *et al*. 2002). Different social rewards have been presented on different days, for example female mice tend to press more for access to a novel female vs. male conspecific when presented on alternating days (Matthews *et al*. 2005), and only female prairie voles pressed more for familiar voles in similar tests (Beery *et al*. 2021). Operant choice tests have typically been used to make comparisons between social and non-social rewards; for instance male Syrian hamsters will overcome a weighted door more often for access to a social chamber with a novel conspecific over an empty chamber (Borland *et al*. 2017). Males of two different strains of mice preferentially lever press for food vs social rewards (Martin *et al*. 2014), while rats press for social rewards vs. food (Hackenberg *et al*. 2021). Social operant studies also routinely employ unfamiliar *strangers* as the social reward for lever pressing. To our knowledge, only one other study has measured social preference in rodents given simultaneous opportunity to access familiar vs. unfamiliar conspecifics (Hackenberg *et al*. 2021). Female rats placed in a two-choice operant apparatus press more for access to a unfamiliar female conspecific (cagemate) over an familiar female conspecific (non-cagemate) (Hackenberg *et al*. 2021). Rats’ novelty preference in an operant social choice test is consistent with their behavior in non-operant social choice tests and peer partner preference tests, in which rats tend to either prefer novelty or lack social preferences, but do not display familiarity preferences (Smith *et al*. 2015; Beery & Shambaugh 2021).

In this study, we developed an operant choice setup to determine whether prairie vole males would work harder for familiar mates than unfamiliar conspecifics, as females were expected to. We assessed whether these voles would exhibit consistent preferences across days for one (clearly preferable) stimulus over another in a non-social choice paradigm (**Fig. 1**) to determine whether learned associations between a particular lever and a particular exposure would occur. We then directly assessed the role of familiarity in social motivation in female and male prairie voles by providing an opportunity for voles to “work” for access to familiar and unfamiliar opposite-sex conspecifics in a two-choice operant apparatus (**Fig. 1**). While prairie voles exhibited consistent preferences in the choice apparatus, there was a striking divergence in social motivation by sex; males pressed a similar amount for both partner and stranger chambers, whereas females preferentially work to access their mate. While males exhibited no overall preferences, individual males differed in their behavior, with some consistently preferring a familiar mate, others preferring a stranger, and yet others exhibiting a lack of preference. This greater heterogeneity among males than females may be related to alternative mating tactics among male prairie voles in the wild. Oxytocin receptor genotype at the intronic locus NT213739 has been associated with oxytocin receptor density and social behavior in prairie voles (King *et al*. 2016; Ahern *et al*. 2021; Beery *et al*. 2021). We replicated a genotype-aggression relationship in males, underscoring oxytocin signaling’s role in the prosocial and selective/antisocial aspects of pairbonding.

**Figure 1.**
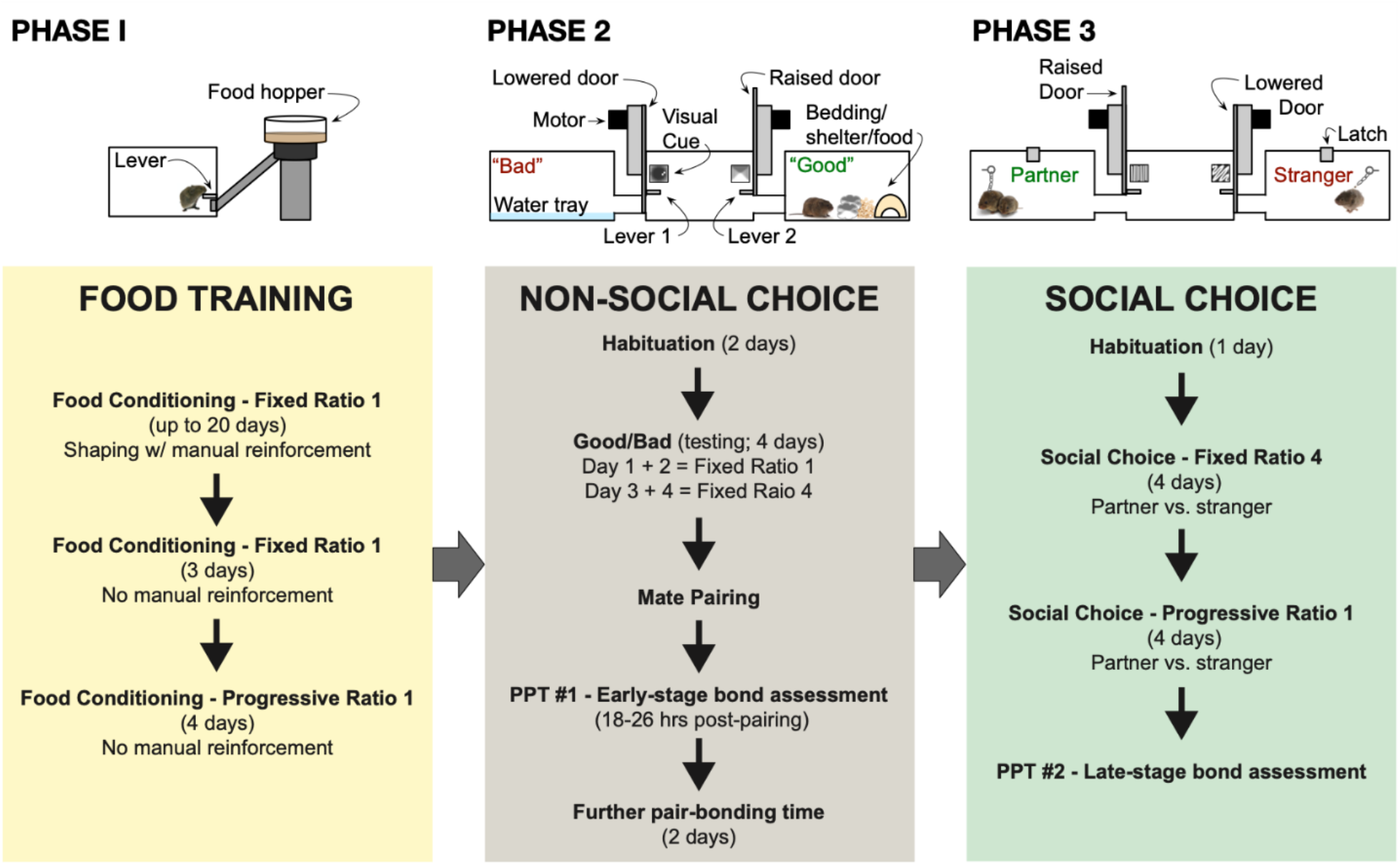
Testing apparatuses and timeline. **Top panel:** Testing apparatus illustration by phase. *Left*: In Phase 1, voles underwent food training within a single-chamber apparatus. In Phases 2 and 3 (*middle* and *right*, respectively), voles were placed in a three-chamber apparatus, wherein the middle operant chamber contained levers that contingently opened a door on either side to a choice chamber containing an environmental (“good” or “bad”; non-social choice) or social opportunity (partner or stranger vole; social choice). **Bottom panel**. Experimental timeline. Apparatus diagrams are shown above the phases with which they were associated.

## Material and Methods

### Animals

Male and female prairie voles (*Microtus ochrogaster*) were bred locally at Smith College, as described in previous experiments (Lee *et al*. 2019; Goodwin *et al*. 2019). Prairie voles were bred in long day lengths (14h light:10h dark) and weaned at 21 days of age. At weaning, subjects were housed in a same-sex pair with an age-matched individual, typically a littermate. Once voles were at least 45 days old, they began the shaping and training protocol. 16 focal animals (7 females; 9 males) completed training and testing with 16 opposite-sex partners (7 females; 9 males) and 30 opposite-sex strangers (14 females; 16 males). All procedures were approved by the Institutional Animal Care and Use Committee of Smith College (ASAF #272) and were conducted in accordance with national guidelines.

### Experimental Timeline

Voles passed through three phases of the study: training, non-social choice operant testing, and social choice operant testing (**Fig. 1**). During Phase 1/food training, focal voles were trained to press a lever for a pellet of food using a single-lever apparatus. Lever pressing behavior was shaped using manual reinforcement, and gradually transitioned to automated reward delivery (described below in *Food training*).

After a focal vole learned the association and motor skill, they transitioned to Phase 2/non-social choice to establish whether voles could learn a stable association between two levers and their rewards, and exhibit a preference. Voles were placed in a two-lever apparatus with chambers on either side of the central chamber. In place of a food reward, lever pressing now resulted in access to two different chambers: one “good” (positive valence environment), and one “bad” (negative valence environment) chamber, described in detail below. After completing food training (∼12 days on average) and the non-social choice “good/bad” protocol (5 days), focal voles were placed in a fresh cage with an opposite-sex vole. 24 hours later, initial bond strength was assessed using a partner preference test (**PPT**). 48 hours following the PPT, voles began phase 3/social choice testing. Focal voles lever pressed for access to two different chambers containing their partner (familiar) and a stranger (unfamiliar) opposite-sex vole. The social choice phase was carried out across eight days. The day after the final social choice operant session, focal voles were administered a second PPT to assess late-stage bond strength. All testing occurred during the light phase. At the conclusion of the study, voles were sacrificed and liver samples were collected for *Oxtr* genotype analysis.

### Gonadectomies & Hormonal Replacement

Prior to the formation of opposite sex pairs, partners of focal voles were rendered infertile in order to prevent pregnancy, as pregnancy can affect the strength of partner relationships (Lewis *et al*. 2017). Female partners of male focal voles underwent tubal ligation to preserve an intact hormonal state. Bilateral incisions were made on the dorsal skin and muscle wall of the voles to access the ovaries. Two knots were placed below each ovary at the top of the uterine horn, sparing the vasculature. The wound was closed using a sterile suture.

Male partners of female focal voles were castrated, then implanted with testosterone (T) capsules to maintain circulating T levels. Testes were accessed by a midline incision, and a tie was placed over the testicular artery to cut off the blood supply. Testes were removed and the muscle wall and skin were closed using sterile suture. Capsules consisted of 4mm of crystalline testosterone (Sigma-Aldrich, St Louis, MO, USA) in silastic tubing (ID 1.98 mm, OD 3.18 mm; Dow Corning, Midland, MO, USA) as in (Costantini *et al*. 2007). Capsules were sealed with silicone, dried, and soaked in saline for 24h prior to implantation. All surgical procedures were performed under isoflurane anesthesia. Voles received 0.05mg/kg buprenorphine and 1.0mg/kg metacam subcutaneously prior to surgery, and again the following day.

### Operant equipment & apparatus

All operant conditioning and testing were carried out in modular test chambers (**Fig. 1**: 30.5 cm x 24.1 cm x 21.0 cm; ENV-307A; Med Associates Inc., St. Albans, VT, USA). For the food training phase, the chamber contained a clicker (ENV-335M) associated with a response lever (ENV-310M), along with a modular pellet dispenser (ENV-203-14P) and receptacle (ENV-303M). For the non-social and social choice phases, the chamber included two response levers and clickers that opened custom guillotine doors (Med Associates Inc.), each adjacent to a lever allowing access to a custom-built chamber (1/4” thick polycarbonate; McMaster-Carr #8574K286) containing an eye-bolt (McMaster-Carr #9489T52). Data were acquired using MED-PC-IV program running custom-coded training protocols.

### Testing schedules

Voles were trained using different operant training schedules as detailed below: fixed-ratio (**FR**), and progressive-ratio (**PR**). FR schedules required a subject to lever press a fixed number of times before a reward was provided (food pellet, chamber access, etc.). For PR, the number of lever presses needed to gain a reward increased proportionally to the number of rewards received. For example, in a PR-1 schedule, one lever press initially yields one reward; however, the vole then needed to press two times to receive the next reward, 3 for the following reward, and so on.

### Food training

In order to motivate voles to learn to associate a lever with reward delivery, voles were placed on food restriction and given food pellets as a reward. Focal voles were weighed two days prior to the start of food restriction. Their average weight across the two days served as a baseline to determine a target weight of 90% relative to their average baseline weight. Two days prior to the start of food training, voles were changed from *ad libitum* food to a diet consisting of two pellets of food (5015; LabDiet, St. Louis, MO, USA) per day maximum. Focal voles undergoing food restriction were weighed daily and their diet was constantly adjusted to prevent their weight from falling below 85% baseline. Focal voles were returned to *ad libitum* food availability after the final day of food training.

Each food training session lasted a maximum of 30 minutes, but was terminated earlier if the vole was inactive. One 20 mg food pellet (Dustless Precision Pellet Rodent Grain Based Diet; Bio-Serv, Flemington, NJ, USA) was placed on the lever before the session began to increase the likelihood a vole would interact with the lever. Initially, a FR-1 schedule was used to train subjects, alongside manual reinforcement by an observer in the room, both of which dispensed a single food pellet. Manual reinforcement was employed if a vole investigated or approached the lever. Voles were transitioned to the next phase (PR-1) for four days after lever pressing at least five times per session on three consecutive FR-1 days without any manual reinforcement (average FR-1 training = ∼12 days).

### Operant non-social choice

Once subjects learned how to lever press, they were transitioned to a three-chamber apparatus for the non-social choice phase (“good/bad”). The goal for this environmental preference test was to verify that voles could learn the association of each lever with its chamber and demonstrate preference for a rewarding chamber over an aversive one through lever presses. Voles underwent daily non-social choice testing for six days.

For the non-social choice tests, the middle chamber contained two levers on opposite sides of the chamber. Each lever was adjacent to a separate door that led to a tube connecting the middle operant (lever pressing) chamber to two other “choice” chambers. Good and bad sides were randomly assigned and counterbalanced across subjects. The good or bad side for each vole remained constant throughout testing (e.g., if the good chamber was on the left side for day 1, it remained on the left side throughout habituation and actual testing days). On the “good” side, voles were presented with a layer of soiled bedding from their home cage, fresh produce (spinach, carrots, and apples), Cheerios, a Shepherd Shack, and a novel object that changed daily (e.g., a metal sports whistle). On the “bad” side, voles were presented with a shallow tray of water (*n* = 12 [7 females; 5 males]).

#### Habituation (days 1 + 2)

On habituation day 1, the levers were concealed, and both choice chambers were empty and accessible for the first 15 minutes. After 15 minutes, voles were shuttled to the operant chamber while the choice chambers were quickly setup. A unique visual cue was placed next to each lever to provide an additional lever/chamber association. Once the setup was complete, the doors were manually reopened for an additional 15 minutes.

On habituation day 2, the levers were concealed, and the doors were propped open, allowing voles free access to both furnished choice chambers for the first 10 minutes. After 10 minutes, voles were shuttled into the operant chamber, the doors were closed, the levers were revealed, and an FR-1 protocol began for 20 minutes. If the vole lever pressed for a given choice chamber, the door was automatically opened for one minute. Voles that remained in the choice chamber after the door closed were immediately shuttled back into the operant chamber.

#### Testing (days 3-6)

The first two days of testing were identical to the latter 20 minutes of habituation day 2, with the entire testing period totaling 30 minutes. For the last two days of testing, the testing schedule was increased to a FR-4.

### Partner preference testing

Partner preference tests were conducted at “early” and “late” stages: 24 hours after pairing with an opposite-sex mate, and at the conclusion of social operant testing, 14 days after pairing. PPTs were carried out as previously described using a three-chambered apparatus connected by tubes (each chamber measured 17 cm × 28 cm × 12.5 cm) (Beery *et al*. 2018; Goodwin *et al*. 2019; Lee *et al*. 2019). Opposite-sex “partner” and “stranger” voles were tethered on opposing sides of the three-chamber apparatus. The untethered focal vole was then placed in the middle chamber and was freely able to explore the entire apparatus for three hours. PPT sessions were video recorded for subsequent offline behavioral analysis. Time in each chamber was scored, as was the time a focal vole spent in physical contact with another subject (“huddling”). The number of aggressive bouts was also noted for each session.

### Operant social choice

The same apparatus was used for social choice as for non-social choice testing. Before a test session was initiated, the focal vole’s partner was tethered to one of the choice sides, and a novel opposite-sex vole (the “stranger”) was tethered to the other choice side. A different (novel) stranger vole was used each for each social choice testing session. As with the choice environment testing, partner/stranger sides were consistent across the entire social choice testing phase. Partner and stranger sides were counterbalanced across subjects such that approximately half of all subjects had the partner on the same side as the former “good” side from the previous phase (non-social choice), whereas the other subjects had the partner on the “bad” side. Novel visual cues were also used for operant social choice training and testing. All social choice testing sessions were video recorded for off-line analysis.

#### Habituation (day 1)

On the first day of social choice, subjects were given 40 minutes to acclimate to the new setup. For the first 10 minutes, the doors were propped open, levers were concealed, and focal voles could freely explore the operant and choice chambers. After 10 minutes, the doors were closed, levers revealed, and an FR-1 protocol was initiated for 30 minutes. *Testing (days 2 – 9)*: On the first four days of testing a FR-4 protocol was used, followed by four days of testing on a PR-1 schedule.

### Behavioral scoring

Videos from testing sessions were analyzed offline by observers using custom Perl scripts (available at https://github.com/orgs/BeeryLab/ and by request). For social tests (social choice; PPT), this yielded the amount of time in resting physical, side-by-side contact with the other voles (“huddling”), the amount of time spent in either choice chamber, the number of entries into either choice chamber, and aggressive bouts. Aggressive bouts were defined as aggressive displays initiated by the focal, including aggressive stances and lunging at the partner or stranger vole.

### *Oxtr* genotyping

DNA used to genotype the NT213739 intronic locus was isolated from frozen liver tissue using the Qiagen DNeasy Kit (Qiagen, #69506), and amplified using forward (5′ - CTCCTATTCAGCCCTCAGAAAC-3′) and reverse (5′ -TGAACCCTTGGTGAGGAAAC-3′) primers, as described in (Ahern *et al*. 2021). The PCR product is a 644 bp amplicon for which BsiHKAI cuts the C-allele to produce bands of 492 bp and 152 bp. Illustra PuRe Taq Ready-to-Go PCR Beads (GE, #27-9557-01) were used with a thermocycler (BioRad) set to 35 cycles (94°C denature, 55°C annealing, 72°C elongation), followed by a 1.5h BsiHKAI restriction digest prior to visualization using a 3% agarose gel (Hoefer, #GR140-500) infused with SYBR green and run for ∼1h at ∼100 V. DNA was analyzed from all 16 focal prairie voles, of which 14 were successfully genotyped (6 females, 8 males). All females were T/T, thus genotype-behavior connections could not be analyzed for this group. Males were either T/T or C/T (“C-carriers”). No C/C individuals were present in the study population; this genotype was also rare in another population and in our prior sample (Ahern *et al*. 2021; Beery *et al*. 2021).

### Statistical analysis

For non-social and social choice tests, lever presses were averaged by subject across the testing phase (four and eight days for non-social and social choice, respectively). For the non-social choice phase, two subjects had more than four days of testing, and the last four days in this phase were averaged instead. Time spent huddling (minutes) was transformed to the percentage of time spent huddling relative to the total available access time. Non-social choice data were analyzed using a 2 × 2 ANOVA (chamber [“good”; “bad”] × sex [male; female]), as there was no *a priori* reason to expect a sex difference for non-social conditions. For the PPT, huddling preference (%) = time huddling (mins) / 180 mins. For the social choice phase, huddling preference (%) = time huddling (mins) / total access time (minutes with that social target’s door raised). Partner preferences were analyzed by 2 × 2 ANOVA (huddling [partner; stranger] × sex). Social choice data were separately analyzed by sex using a 2 × 2 ANOVA (chamber [partner; stranger] × schedule [d1-4/FR; d5-8/PR]). Individual paired *t*-tests were run across social choice data for lever presses × chamber (partner; stranger) to determine chamber preference. Aggressive bouts data were analyzed using a 2 × 2 ANOVA (average bouts across the eight days of social choice × chamber). For aggression correlation analyses, aggressive bouts were scaled relative to the number of door openings (aggressive bouts / total access opportunities). Statistical analyses were performed using RStudio version 1.4.1103 (RStudio Team 2020) running R version 3.5.2 (R Core Team 2018) using the following packages: *tidyverse* (Wickham *et al*. 2019); and *stats* (aov; TukeyHSD [adjusted for multiple comparisons]) (R Core Team 2018). Comparisons of behavior by *Oxtr* genotype were conducted by Welch’s *t*-test assuming unequal variances. Two female subjects were excluded from all analyses due to low operant activity (< 20 lever presses in total across all eight days of the social phase). Four PPT video recordings were incomplete and not used, and one male was excluded from PPT1 analysis due to huddling less than five minutes in total (as in Anacker *et al*. 2016). The data that support the findings of this study are available in the Open Science Framework under the project title “Sex differences in the reward value of familiar mates in prairie voles” at https://osf.io/wmjfs/.

## Results

### Non-social (environmental) preferences

Non-social operant choice tests were used to establish whether voles would consistently learn to associate specific levers with different stimuli, and lever press more for the preferred stimulus. A two-way ANOVA (sex [male; female] * chamber [good; bad]) revealed that voles lever pressed more for access to the “good” chamber (*F*_(1, 10)_ = 18.627, *p* = 0.002; **Fig. 2**). There was no significant effect of sex (*p* = 0.392) nor a sex * chamber interaction (*p* = 0.548). This indicates that both male and female voles are able to reliably distinguish between chambers and the corresponding levers that provides access. Thus, any lack of preference would not indicate lack of learning ability in the test.

**Figure 2.**
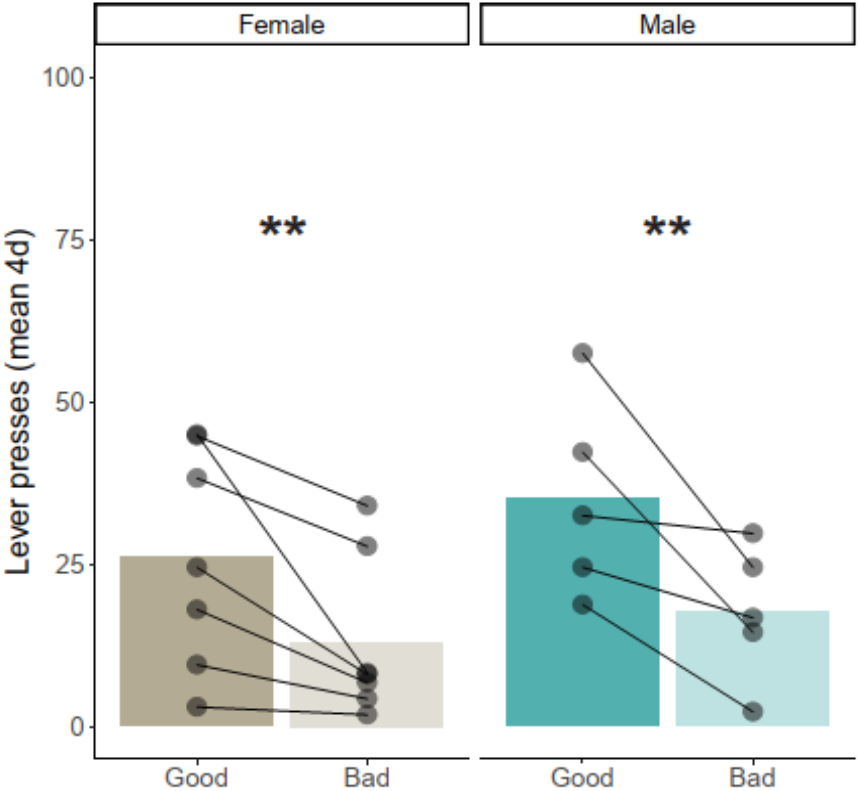
Prairie voles can distinguish between negative (“bad”) and positive (“good) valence environments, and demonstrate preference for the latter. Males and females alike selectively lever pressed for access to the “good” environment of the “bad” one during the non-social choice phase. Dots represent the mean number of lever presses over 4-days of testing. Bars represent group means. Asterisks indicate significant environment preference within sex. ** *p* < 0.01.

### Social motivation

#### Lever pressing

To evaluate whether voles spent more time working towards access for their partner or a stranger, we compared the number of lever presses for each sex across the social choice phase. Males pressed similar amounts for the stranger and partner (*F*_(1,8)_ = 0.66, *p* = 0.44; **Fig 3A**), irrespective of schedule (*F*_(1,8)_ = 4.64, *p* = 0.063) or interaction between chamber and schedule (*F*_(1,8)_ = 1.74, *p* = 0.22). In contrast, females lever-pressed significantly more for their partner over a stranger vole (*F*_(1,4)_ = 15.37, *p* = 0.017; **Fig 3A**), with no effect of schedule (*F*_(1,4)_ = 5.67, *p* = 0.49), nor an interaction between schedule and chamber (*F*_(1,4)_ = 0.57, *p* = 0.49).

**Figure 3.**
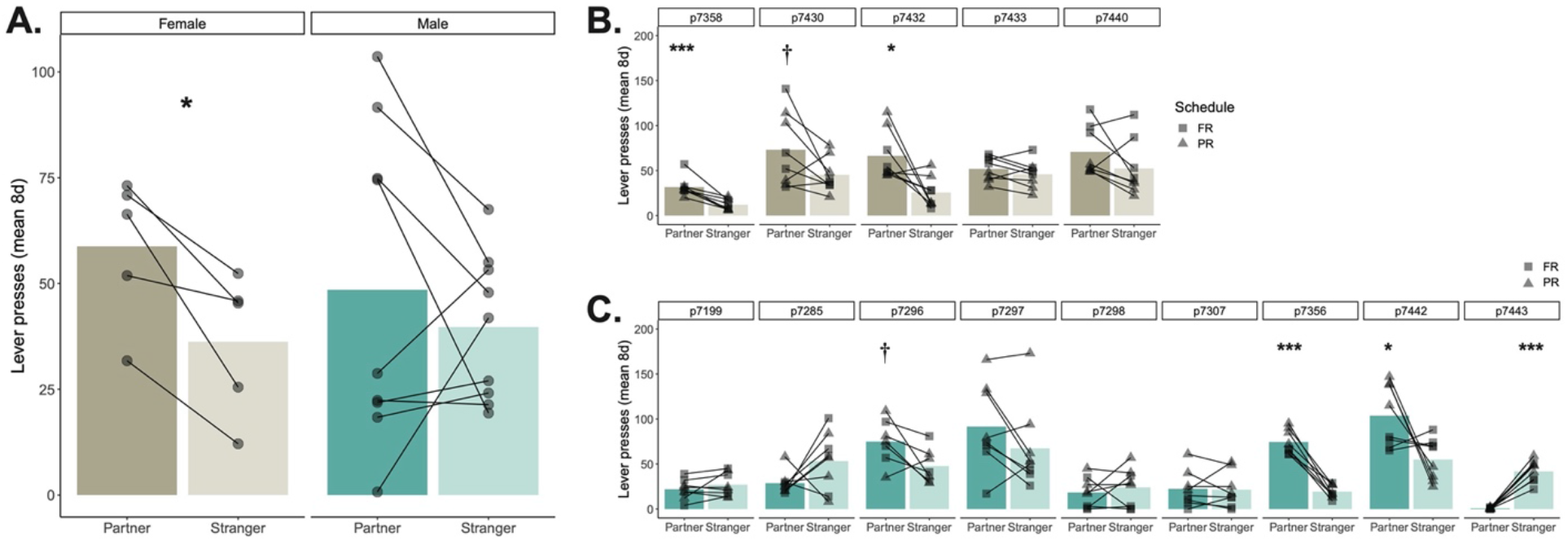
Sex-specific lever pressing rates for access to a familiar versus unfamiliar opposite-sex conspecific. **A**. Female prairie voles lever pressed more on average for their mate than for an unfamiliar opposite-sex vole, whereas males pressed at similar rates for both their mate and the novel conspecific. **B. & C**. Individual lever pressing patterns across the 8 days of testing. Numbers at the top are vole identifiers. Paired *t-*tests for lever pressing within subjects reveal consistent individual differences in preference patterns across testing days, especially in males **(C.)**. *** *p* < 0.001; ** *p* < 0.01; * *p* < 0.05; † p < 0.1.

As there was notable individual variation in lever pressing preference, particularly among males, we followed up group analyses with individual analyses. When daily pressing of individual subjects was examined across the social choice phase, three of five females significantly or marginally (*p* < 0.1) preferred their partners, while the remaining two pressed more for their partner on average, but with no significant difference (**Fig. 3B**). Among males, three of nine significantly or marginally preferred their partners, one significantly preferred the stranger female, and the remaining five exhibited no consistent preferences (**Fig. 3C**).

### Huddling

In addition to analyzing lever pressing, we also explored the amount of time focal voles spent huddling with their partner or a stranger. During PPT1, both males and females significantly preferred huddling with partner over the stranger (**Fig. 4A**; huddling stimulus (partner; stranger): *F*_(1,9)_ = 138.16, *p* < 0.001; sex: *F*_(1,9)_ = 1.72, *p* = 0.22; sex * stimulus: *F*_(1,10)_ = 1.94, *p* = 0.20). These strong partner preferences in both male and female huddling times are consistent with prior studies.

**Figure 4.**
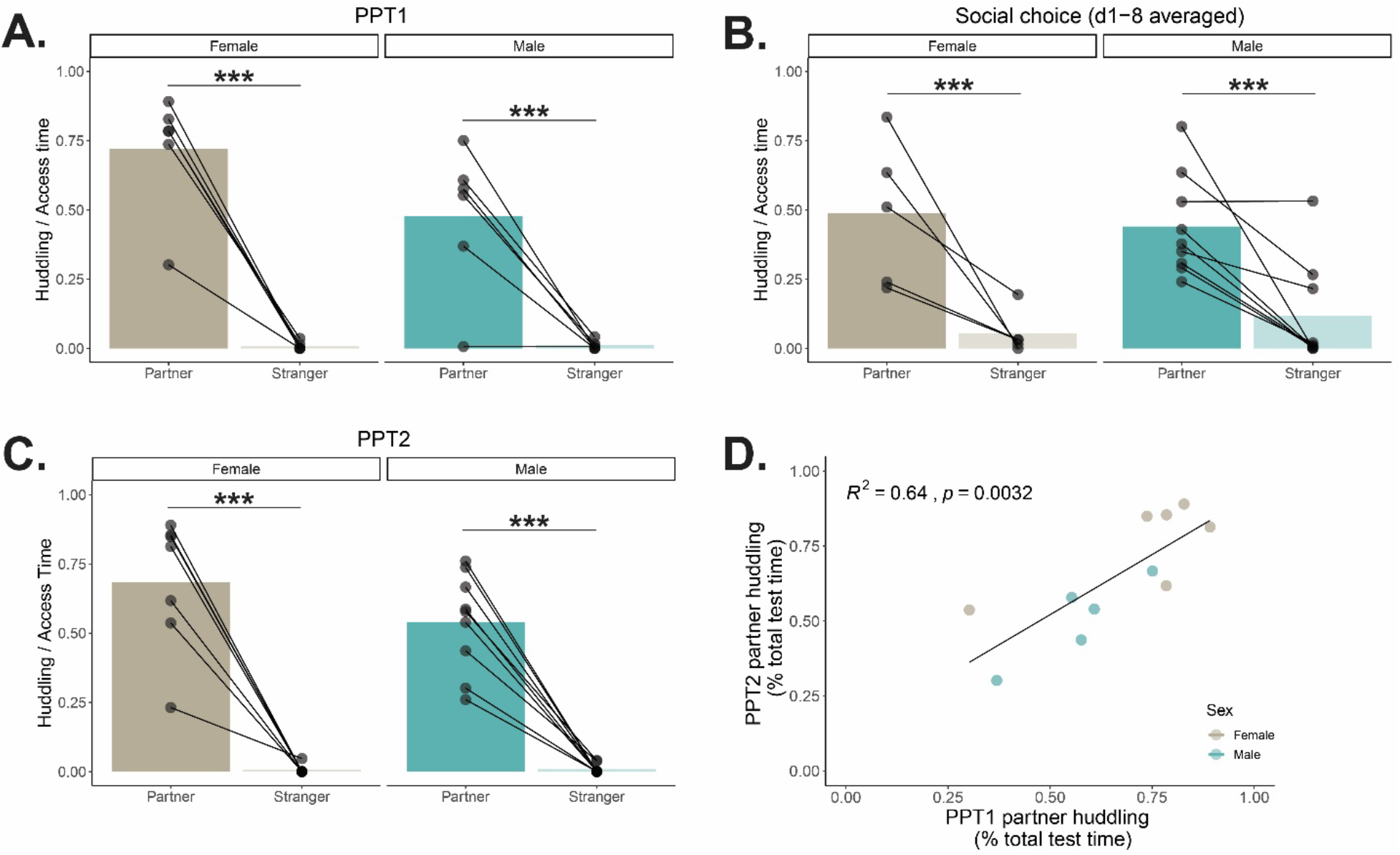
Prairie voles prefer huddling with their mate irrespective of sex. Both males and females exhibited strong partner preferences in 24 hours after pairing (PPT1; **A**.), across the operant social choice testing days (**B**.), and in PPT2, 2 weeks after pairing (**C**.). **D**. Partner huddling was consistent within individuals across PPT sessions. Males and females exhibited the same patterns, and this correlation was individually significant in males. *** *p* < 0.001.

Similarly, when huddling was assessed across the eight days of social choice, both male and female voles spent more time huddling with their partner over a stranger vole relative to total available time (**Fig. 4B**; stimulus type: *F*_(1,12)_ = 30.651, *p* < 0.001; sex: *F*_(1,12)_ = 0.009, *p* = 0.92; sex * stimulus vole: *F*_(1,12)_ = 0.66, *p* = 0.43). Analyzing partner or stranger huddling relative to either the total available time for a specific stimulus vole or total test time yielded similar results. We also investigated whether subjects’ pair-bond preference shifted following operant social choice during a late-stage pair-bond assessment (**PPT2**; see **Fig. 1B**). Just as before in PPT1 and during operant social choice, focal subjects spent more time huddling with their partner over a stranger vole, independent of sex (**Fig. 4C**; stimulus vole: *F*_(1, 14)_ = 121.93, *p* < 0.001; sex: *F*_(1,14)_= 1.99, *p* = 0.18; sex * stimulus type: *F*_(1,14)_ = 1.80, *p* = 0.20).

Finally, we assessed whether the preference for the partner vole was consistent within subjects across the experiment. Because there were no sex differences in partner preference, males and females were analyzed together, and displayed a significant positive correlation for PPT1 to PPT2 huddling time with their partner (*r*^2^ = 0.64; *p* = 0.003; **Fig. 4D**). Sub-analysis by sex confirmed that male voles retained a significant positive correlation for huddle time with their partner from PPT1 to PPT2 (*r*^2^ = 0.82; *p* = 0.03), while females showed a positive correlation that approached significance (*r*^2^ = 0.55; *p* = 0.09; **Fig. 4D**). Females typically huddled more with their partner than males do as evidenced by clustering of females at the top right of Fig 4D (as in Brusman et.al., submitted to the same issue). Due to a lack of variability in individual data (i.e. low stranger huddling), we were unable to assess similar correlations for stranger huddling during PPT1 and PPT2.

### Aggression

Both males (*F*_(1,8)_ = 24.90, *p* = 0.001) and females (*F*_(1,4)_ = 22.15, *p* = 0.009) were more aggressive towards the stranger (S) vole compared to their partners (P) during operant social choice (males: 2.01 ± 0.40 S bouts, 0.09 ± 0.07 P bouts; females: 1.30 ± 0.28 S bouts and 0.10 ± 0.10 P bouts). Because aggressive behavior can be rewarding (e.g. in socially dominant male mice, (Falkner *et al*. 2016; Golden *et al*. 2019), we asked whether stranger-directed aggression was correlated with lever pressing activity. There were no significant correlations between lever pressing and aggression relative to access time, and if anything there was the opposite relationship, with females that pressed more for the stranger exhibiting less likelihood of aggression (*R*^2^ = 0.09, p = 0.058) and no relationship in males (*R*^2^ = 0.03, *p* = 0.16). Opportunities for aggression were not a motivating factor for access to unfamiliar females’ chambers.

We previously discovered a robust connection between aggressive behavior towards unfamiliar voles and *Oxtr* genotype at the intronic locus NT213739 in males (Beery *et al*. 2021); this SNP has been associated with both oxytocin receptor binding density within the nucleus accumbens, and preference behaviors (King *et al*. 2016; Ahern *et al*. 2021). The present sample skewed strongly toward T/T individuals (6 of 6 females, and 6 of 8 males that were successfully genotyped). Even with low sample diversity and size, we replicated the association between genotype and aggressive behavior in males, with T/T individuals exhibiting more bouts of aggression towards strangers relative to access time than C/T individuals (mean 0.45 ± 0.1 versus 0.14 ± .09 bouts/minute of access; *t*(4.2) = 2.23; *p* = 0.04; one-tailed). Because aggression/access time is a comparable variable across the two social operant studies conducted in our lab, we also computed the omnibus relationship across the present study and Beery et al. 2021. In the combined sample of 16 T/T males and 6 C/T males, aggression was again over three times higher in T/T males (mean .57 ± 0.1 bouts/min versus .17 ± .06 bouts/min of access time; *t*(19.9)=3.2; *p*<0.005).

## Discussion

Social contact with mates is behaviorally rewarding for prairie voles, and there have been some indications of sex differences in reward value (Ulloa *et al*. 2018; Goodwin *et al*. 2019; Beery *et al*. 2021). The present study extends these findings, demonstrating that even when faced with a direct choice between a partner and a stranger, males do not consistently work harder to access their partner, unlike female prairie voles. This sex difference reveals a striking disconnect between social motivation and partner preference, as both male and female voles exhibited robust preferences for their mates over opposite-sex strangers in partner preference tests, as well as in huddling/access time within the operant behavioral tests. Males also exhibited considerable individual variation in behavior, potentially reflective of the diversity of mating tactics and behavioral strategies exhibited in the wild (Shuster *et al*. 2019; Madrid *et al*. 2020).

### Sex differences

Monogamous species often exhibit fewer overt sex differences than promiscuous species, both physically, in terms of similar appearance, and behaviorally, for instance engaging in biparental care and forming partner preferences for a pair-bonded mate. Such behavioral similarity between the sexes may, however, arise from sex-specific mechanisms that compensate for differences in gonadal hormone exposure (De Vries & Boyle 1998; De Vries 2004). These so-called “latent” sex differences (also referred to as mechanistic, convergent, or divergent sex differences) often manifest as small but repeatable sex differences in behavior that can mask larger sex differences that may be revealed once underlying mechanisms are probed (Beltz et al., 2019; for sex-specific pain processing as an example, see Mogil, 2018).

Early in the study of prairie vole pair-bonding, oxytocin was deemed more important for partnership formation in females than in males, and vasopressinergic pathways became the focus of studies on male prairie voles (Winslow *et al*. 1993). In subsequent years, most studies have focused on neuropeptide, dopaminergic, and opioid signaling pathways in one sex at a time, although studies in one sex and then the other reveal potentially similar roles of some of these pathways (Johnson *et al*. 2016; Resendez *et al*. 2016). To identify or quantify sex differences, it is critical to include males and females in the same study. While this practice is increasingly common, analysis of subjects by sex is still far from the norm (Beery & Zucker 2011; Woitowich *et al*. 2020). When direct comparisons of males and females have been made in prairie voles, it has revealed sexual dimorphisms in the effects of stress on social bond formation (DeVries *et al*. 1996), sex-specific effects of pair bonding on kappa opioid receptor densities, and sex differences in the relative importance of familiarity in social reward in (present study and (Beery *et al*. 2021)). A related study also found that the behavioral factors that contribute to partner preference in male and female voles differ—females increase partner-directed huddle while males decrease novel-directed huddle as pair bonds mature. The same study also used an operant social task similar to the one presented here and found that females but not males exhibited differences in effort to access a mate vs. stranger (Brusman et al, submitted to this issue of GBB). Together, these results all point to sex differences in reward and motivation as they relate to the display of key pair bonding behaviors.

The decoupling of social reward and social preference (i.e. selective huddling behavior in males) indicates that stranger females have rewarding properties not captured by their desirability as a target of social huddling. Extra-pair copulation opportunities are one obvious potential source of reward. Aggression may provide another possible source of reward from stranger contact (Falkner *et al*. 2016; Golden *et al*. 2017), although this is unlikely to explain our results. While prairie voles display relatively high levels of stranger-directed aggression (Lee et al., 2019) which may be an important reinforcer of pair bonds (Aragona *et al*. 2006; Young *et al*. 2011), in the present study, there was no relationship between stranger-directed aggression and lever pressing effort, and there was a non-significant negative relationship between stranger-directed aggression and lever pressing for the stranger, suggesting that aggression was not a particular motivator for males in accessing strangers.

### Heterogeneity in male behavior

While all voles exhibited strong partner preferences, there was extensive individual variation in lever pressing preferences for the partner vs. stranger. Males, in particular, displayed distinct but consistent classes of social preferences: some pressed significantly more for the partner, others pressed more for the stranger, whereas others pressed similar amounts for both conspecifics. Females were less variable, with some females exhibiting significant preferences for their partner vs. a stranger over the 8 day testing interval, and others not exhibiting significant preferences, but all females pressed more for their partner than for the stranger on average.

Increased heterogeneity in social motivation in males in consistent with inter-male variation in mating strategies in the field. While most prairie voles are socially monogamous (and exhibit a “resident” strategy for mate partnerships), a notable percentage of both male and female prairie voles (25 – 40%) are non-monogamous (sometimes referred to as “wanderers”) (Madrid *et al*. 2020). Some of the variation in mating tactics may be due to environmental conditions (Mabry *et al*. 2011; Madrid *et al*. 2020). Our finding extends this distribution of individual variability in mating tactics and quantifies it using an operant paradigm.

### Oxtr polymorphism and social behavior

Prior work suggests that a SNP in the intron of the prairie vole *Oxtr* gene NT213739 contributes to individual differences in striatal OXTR protein levels and attachment behavior (King *et al*. 2016; Ahern *et al*. 2021). Genetic diversity in the sample used for the present study was too low to analyze female data, or to discover new patterns in the male data. Nonetheless, the male data replicated our prior finding that C allele carriers (C/T) were significantly less aggressive than T/T homozygous individuals (Beery et al., 2021), and added robustness to the prior data set. Partnership formation in prairie voles occurs alongside increases in stranger-directed aggression (Winslow *et al*. 1993; Insel *et al*. 1995; Wang *et al*. 1997; Aragona *et al*. 2006), thus these physiological and behavioral outcomes are concordant.

In sum, by developing direct measures of partner-directed motivation in a choice context, we quantified a distinct behavioral component implicated in pair bonding. In doing so, we identified robust sex differences in the role of reward in preference behaviors. This advance is critical for subsequent investigation of the neural and genetic systems that contribute to pair bond motivation in males and females, as well as for parsing a well-delineated example of “latent” sex differences.

## Acknowledgements

We are grateful to Jessie Chen, Juliane Donahue Bombosch, Maddie Lerner, and Katerina Rusa for assistance running and scoring behavioral tests, and to Anna Pintchouk, Isobel Abbot, and Xueling Zhang for assistance running tests. Nikki Lee assisted with prairie vole colony maintenance and general know-how. Sarah Lopez and Dale Renfrow (Smith Center for Design and Fabrication) designed and constructed the social testing chambers. We thank the staff of the Smith College Animal Care Facility for animal care and colony maintenance. This research was supported by the National Institute of Mental Health of the National Institutes of Health under Award Number R15MH113085 (to A.K.B.) and DP2OD026143 (to Z.R.D.)

